# Tissue heterogeneity is prevalent in gene expression studies

**DOI:** 10.1101/2020.12.02.407809

**Authors:** Gregor Sturm, Markus List, Jitao David Zhang

**Affiliations:** Biocenter, Institute of Bioinformatics, Medical University of Innsbruck, 6020 Innsbruck, Austria; Chair of Experimental Bioinformatics, TUM School of Life Sciences Weihenstephan, Technical University of Munich, 85354 Freising, Germany; Roche Pharma Research and Early Development, Pharmaceutical Sciences, Roche Innovation Center Basel, F. Hoffmann-La Roche Ltd, Grenzacherstrasse 124, 4070 Basel, Switzerland

## Abstract

**Background:** Lack of reproducibility in gene expression studies has recently attracted much attention in and beyond the biomedical research community. Previous efforts have identified many underlying factors, such as batch effects and incorrect sample annotations. Recently, *tissue heterogeneity*, a consequence of unintended profiling of cells of other origins than the tissue of interest, was proposed as a source of variance that exacerbates irreproducibility and is commonly ignored.

**Results:** Here, we systematically analyzed 2,692 publicly available gene expression datasets including 78,332 samples for tissue heterogeneity. We found a prevalence of tissue heterogeneity in gene expression data that affects on average 5-15% of the samples, depending on the tissue type. We distinguish cases of severe heterogeneity, which may be caused by mistakes in annotation or sample handling, from cases of moderate heterogeneity, which are more likely caused by tissue infiltration or sample contamination.

**Conclusions:** Tissue heterogeneity is a widespread issue in publicly available gene expression datasets and thus an important source of variance that should not be ignored. We advocate the application of quality control methods such as *BioQC* to detect tissue heterogeneity prior to mining or analysing gene expression data.

## Background

The genome-research community has witnessed the exponential growth of gene expression studies in the last two decades, first with microarray^1^ and nowadays with RNA-seq datasets^2^. Both the huge volume of data and wide coverage of biological samples in diverse contexts, such as genetic perturbation, disease progression, pharmaceutical intervention, *etc*., make publicly available gene expression studies an important resource for drug-discovery research. Systematic mining of of existing data and interrogation of new data can reveal molecular foundations of pathology and disease^3^, identify novel therapeutic targets^4^, enable preclinical screening tools for preclinical drug safety^5,6^, highlight mode-of-action of drug candidates ^7^, allow data-driven prioritisation of drug screening hits^8^, and predict response and stratify patients^9^. In short, drug discovery benefits from both consuming and contributing to gene expression studies.

However, the power of gene expression studies in translating molecular biology into medicine is impeded by a lack of reproducibility^10,11^. Well known causes of irreproducibility include batch effects, variation of biological samples, profiling protocols, or data analysis procedures, as well as mistakes in sample handling or annotation, and in rare cases intentional data manipulation. While several studies have scrutinized publicly available gene expression datasets, and demonstrated the prevalence of *e*.*g*. batch effects^12^, and sample mishandling or misannotation^13^, few studies address the prevalence of *tissue heterogeneity, i*.*e*. the unintended profiling of cells of other origins than the tissue of interest^14,15^. Tissue heterogeneity can be caused by intrinsic characteristics of the sample to be profiled, such as the tumor microenvironment or immune cell infiltration into solid organs, or by extrinsic factors such as imperfect dissection or contamination of samples. Ignoring tissue heterogeneity reduces statistical power of data analysis and can, in the worst case, invalidate the conclusions of a study. In particular in oncology, this is a well recognized problem that is commonly addressed by estimating tumor purity^16^. Alternatively, cell type heterogeneity can be leveraged as a source of information in immune cell deconvolution to inform about the state of the tumor microenvironment and to guide immunotherapy^17^. Beyond tumor samples, Nieuwenhuis et al. identified a cluster of prostate-specific genes that were expressed in tissues other than prostate in GTEx, but also in other datasets, highlighting that tissue contamination is an important issue found in important reference datasets commonly used by the community^15^. To our knowledge, a systematic analysis of cross-tissue contamination in which data sets are systematically tested for contamination with other tissues than the tissue of interest is missing. Here, we systematically screened 2,692 datasets from the two largest public microarray and RNA-seq gene expression repositories, Gene Expression Omnibus (GEO)^18^ and ARCHS4^19^, respectively, for tissue heterogeneity using the R package *BioQC*^*14*^. We found that, independent of the technological platform but depending on the tissue type, between 5-15% of samples suffered were affected by tissue heterogeneity with on average 1.6% suffering from severe heterogeneity issues (FDR threshold=0.01). Our results, thus, highlight the importance of considering tissue heterogeneity as a confounder in transcriptome data analysis.

## Results and Discussion

We evaluated the enrichment of 120 different *query signatures* from the R package *BioQC* in a selection of well annotated microarray studies in the GEO^18^ and ARCHS4^19^ repositories (2,692 studies, 78,332 samples, Figure 1A-B). These query signatures are tissue-sensitive, *i*.*e*. they recognize their target tissue with few false negatives, but often not tissue-specific, *i*.*e*. they report false positives due to the expression of the signature genes in other, physiologically similar tissues. To account for this, we created a set of nine *reference signatures* using GTEx data^20^ and validated them using the GNF MouseAtlas V3 dataset^21^ to show that they are robust even across species (Figure 1C). To identify samples affected by heterogeneity, we apply a two-step approach (Figure 1D): (1) the enrichment score of the query signature needs to exceed an absolute cutoff; (2) for each query signature, we fit a linear model of the query signature score against the reference signature score, where the reference signature matches the sample’s tissue. If the query signature score cannot be explained by the linear model, we consider the sample heterogeneous. The cutoffs were chosen such that the overall FDR of the two-step testing procedure is 0.01.

**Figure 1.**
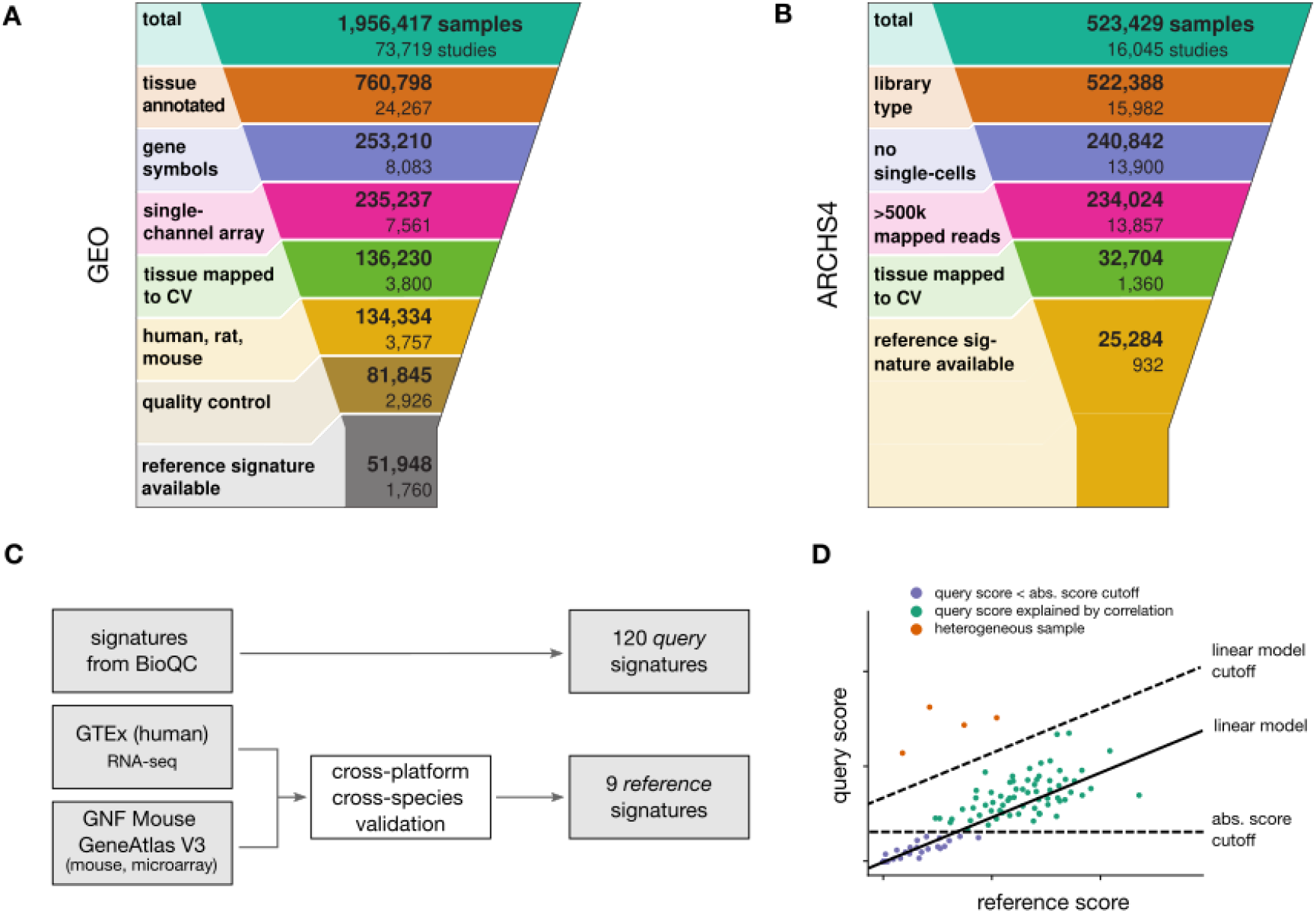
**(A-B)** Selection of gene expression studies from **(A)** GEO and **(B)** ARCHS4. **(C)** We defined two sets of tissue signatures for the experiment: (1) we obtained 120 tissue *query* signatures from the *BioQC* package and (2) generated 9 high-quality *reference* signatures from the GTEx and GNF Mouse GeneAtlas V3 datasets. **(D)** Schematic illustration of the two-step approach to call heterogeneous samples. Since query signatures may be imperfect and correlated with the sample’s tissue of origin, we use a linear model to compare the query against a robust reference signature. Abbreviations: CV, controlled vocabulary.

We further distinguish between severe and moderate tissue heterogeneity. Empirically, we define moderate heterogeneity as samples that are significantly enriched for a signature that we do not expect to be present, and severe heterogeneity as samples in which, in addition, the expected signature of the annotated tissue is not detected. While severe heterogeneity often suggests mistakes in sample handling and annotation, moderate heterogeneity suggests contamination or infiltration of cells of the blood and immune system.

We found moderate tissue heterogeneity in about 6.6% of all samples and severe heterogeneity in 1.6% of samples. The proportion of samples affected by moderate heterogeneity varies by the organ and tissue being profiled, with skin (14%) and pancreas (14%) samples affected most frequently and brain samples affected least frequently (<5%) (Figure 2A). Results were comparable for RNA-sequencing and microarray, corroborating that the issue of sample heterogeneity is not platform-dependent.

**Figure 2:**
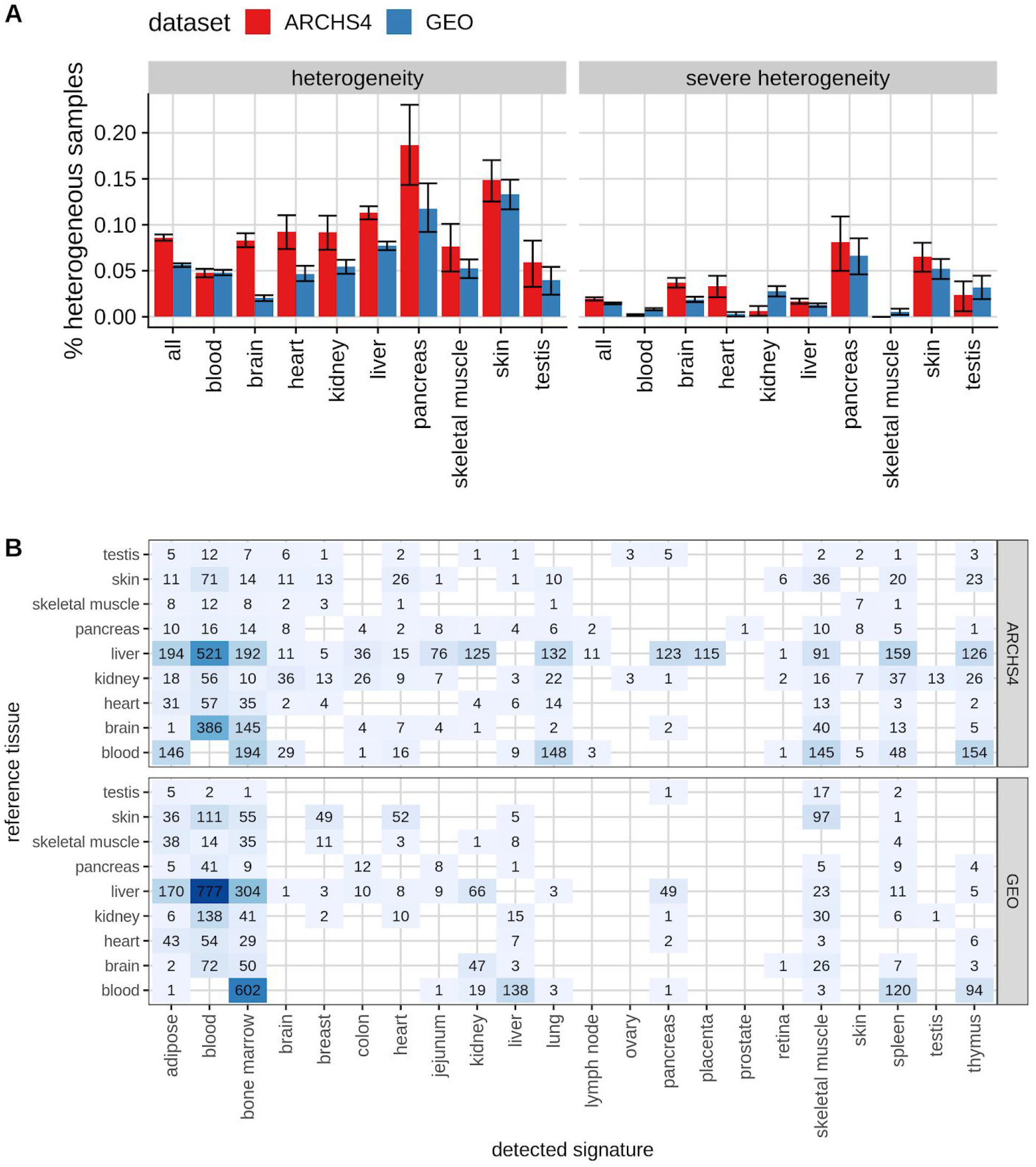
Tissue heterogeneity in gene expression studies from GEO and ARCHS4. **(A)** Fraction of heterogeneous samples per tissue. Error-bars show 95%-confidence intervals derived by bootstrapping. **(B)** Tissue confusion matrix with absolute counts. Reference tissue refers to the annotated tissue, detected signature to other tissue signatures that were detected in these samples by BioQC.

A closer investigation of the source of tissue heterogeneity reveals additional insights (Figure 2B). For instance, enrichment of blood signatures in other tissues and organs is one of the most frequent forms of severe heterogeneity which can be caused by an increased inflow and/or decreased outflow of blood which sums as a net increase of blood volume, or the activation and proliferation of tissue-resident leukocytes, for instance. Tissue heterogeneity of proximal tissues could be explained by imperfect separation of nearby organs. For example, the liver and pancreas are proximal organs connected by the common bile duct, which may explain why many cases of tissue heterogeneity in pancreatic tissue are caused by liver-specific tissue signatures. However, tissue heterogeneity involving distal solid tissues also occurs and highlights possible issues with contamination during sample preparation. Considering that the latter two aspects represent technical biases, such samples should be excluded in analysis to increase statistical robustness and to avoid arriving at erroneous conclusions.

A limitation we encountered in this study were normalized expression profiles in the GEO repository. As *BioQC* depends on the ranking of genes within a sample, studies that performed a per-gene normalization could not be evaluated. We, therefore, advocate the upload of the raw data to gene expression repositories as only the unmodified gene expression data can be used with *BioQC* or other quality control tools.

We also note that the issue of tissue heterogeneity is specific to bulk RNA-sequencing data and does not affect single-cell RNA-seq studies, as contaminating cells form an independent cluster of cells which can either be ignored or incorporated in data analysis. In fact, single-cell RNA-seq offers the chance to study biological sources of tissue heterogeneity at a previously unimaginable depth. However, due to its lower cost, the majority of expression profiles will still be sequenced in bulk in the foreseeable future. Hence, identifying samples affected by tissue heterogeneity with tools such as *BioQC* will remain an important aspect of data analysis and should be incorporated in standard gene expression analysis pipelines.

## Conclusions

Tissue heterogeneity is prevalent in publicly available gene expression data and occurs independent of the technological platform. Tissue heterogeneity affects on average 6.6% of samples with differences between tissues ranging from 5% (brain) to 14% (skin, pancreas). While most cases of moderate heterogeneity are caused by blood cells, enrichment with signatures from proximal and distal solid tissues highlights contaminated samples that should be excluded from analysis. Similarly, samples with severe heterogeneity (on average 1.6%) should be excluded. This is of particular importance in systems medicine studies, where tissue-specific signals can mask disease-specific signals, thus preventing the successful detection of disease mechanisms, patient stratification, or drug target identification and validation. To avoid this, we advocate the routine use of methods such as *BioQC* that assess tissue heterogeneity in transcriptome analysis.

## Methods

### Compilation and cross-validation of tissue signatures

*BioQC* provides 155 sets of tissue-enriched genes (tissue signatures hereafter) from four large-scale tissue gene expression datasets^14^. Even though the authors have shown that the signatures are biologically meaningful, they did not validate them using an independent dataset. Since the reliability of signatures is crucial for this study, we developed an open-source software package, *pygenesig*, which facilitates the creation and validation of tissue signatures. We applied *pygenesig* to transcriptomics data from the GTEx project^20^ (v6) which contains 11,984 samples from 32 tissues and validated the resulting signatures on the GNF Mouse Gene Atlas V3^21^. We identified a set of 9 “reference” tissue signatures, that reliably identify their tissue of origin, regardless of experimental platform and species. The process of signature generation and validation is outlined in Figure 1C and detailed in supplementary section 2.

### Gene expression data corpus

We retrieved annotation and gene expression data from GEO on 2016-12-07 using *GEOmetadb*^*22*^ and *GEOquery*^*23*^. We downloaded consistently processed RNA-seq gene expression data including annotations as RData objects from the ARCHS4 project website (version 8.0)^19^. Data filtering and quality control is summarised in Figure 1A-B and described in detail in supplementary section 3.

Tissue annotations in GEO and ARCHS4 are inconsistent. We, therefore, manually mapped tissue descriptions to a controlled vocabulary, and assigned 120 of the 155 signatures provided by *BioQC* and the 9 reference signatures to their corresponding tissues (supplementary table 1).

### Detecting tissue heterogeneity with BioQC in the corpus

*BioQC* performs a Wilcoxon-Mann-Whitney statistical test for enrichment of a certain signature on a per-sample basis. We ran *BioQC* on all samples from GEO and ARCHS4 using both the nine reference signatures and 120 signatures provided by *BioQC*, which yielded 10,031,456 (sample, signature, p-value) pairs. As signatures can be correlated (*e*.*g*. because they describe physiologically related tissues), we apply the following procedure to identify heterogeneous samples: A given sample *s* annotated as tissue *t* is tested for enrichment with the query-signature *k*_query_ resulting in a p-value *p*_query_. Let *k*_ref_ be the reference signature associated with tissue *t* and *p*_ref_ the p-value of testing *s* for enrichment of *k*_ref_. Let τ be the false-discovery-rate (FDR) threshold. (1) If the Benjamini-Hochberg (BH)-adjusted *p*_query_ ≥ τ, we label *s* as not heterogeneous, else continue. (2) We fit a robust linear model using `rlm` from the R package *MASS* of |log_10_(*p*_query_)| against |log_10_(*p*_ref_)| for all samples annotated as *t*. We assume that the residuals *R* of the linear model follow a normal distribution *R* ∼ *N* (0, σ^2^), where σ is the standard deviation of the residuals. (3) We extract the residual *r* corresponding to sample *s*. We calculate the p-value *p*_corr_ = 1 - CDF_*R*_(*r*), where CDF_*R*_ is the cumulative density function of *N* (0, σ^2^). (4) If the BH-adjusted *p*_corr_ < τ, we reject the hypothesis that *k*_query_ is enriched only due to correlation and label the sample as heterogeneous. We choose τ such that the overall FDR of the two-step testing procedure equals 0.01. This process is illustrated in figure 1D and supplementary section 4.

Finally, we computed the fraction of heterogeneous samples by dividing the number of samples that have at least one signature passing the above criteria by the total number of samples per tissue. Confidence intervals have been derived by bootstrapping using the R package *boot*.

### Other software

We implemented and documented the analysis using R bookdown^24^. The analysis is wrapped into a reproducible pipeline built on Snakemake^25^.

## Declarations

### Ethics approval and consent to participate

Not applicable.

### Consent for publication

Not applicable.

### Availability of data and material

The reference signatures and raw results table including all accession numbers tested is available from https://doi.org/10.5281/zenodo.4298774. The source code to reproduce the analysis can be found at https://github.com/grst/bioqc_geo. *Pygenesig* is available from https://github.com/grst/pygenesig. *BioQC* is available from https://accio.github.io/BioQC/.

### Authors’ contributions

GS and JDZ conceived the study. GS curated microarray datasets and performed analysis. GS and JDZ collected RNA-sequencing datasets and performed analysis. GS, ML and JDZ wrote the manuscript and approved its content.

## Acknowledgements

We would like to thank the members of the bioinformatics group at Roche for inspiring discussions. Especially we thank Laura Badi for her continuous work to improve the *BioQC* method, Klas Hatje, Iakov Davydov, and Roland Schmucki for testing *BioQC* and providing valuable feedback, Martin Ebeling and Manfred Kansy for suggesting the case study and supporting the work. JDZ wishes to dedicate this study to Clemens Broger, in memory of his inspiring and loving style of working with young people.

## Supplementary Material

- Supplementary Information
- Supplementary Table 1: Tissue mapping table

## References

1. Baker M. Gene data to hit milestone. Nature 2012;487:282–283.

2. Stark R, Grzelak M, Hadfield J. RNA sequencing: the teenage years. Nat Rev Genet 2019;20:631–656.

3. Xu J, Acharya S, Sahin O, et al. 14-3-3ζ turns TGF-β’s function from tumor suppressor to metastasis promoter in breast cancer by contextual changes of Smad partners from p53 to Gli2. Cancer Cell 2015;27:177–192.

4. Moisan A, Lee Y-K, Zhang JD, et al. White-to-brown metabolic conversion of human adipocytes by JAK inhibition. Nat Cell Biol 2015;17:57–67.

5. Moisan A, Gubler M, Zhang JD, et al. Inhibition of EGF Uptake by Nephrotoxic Antisense Drugs In Vitro and Implications for Preclinical Safety Profiling. Mol Ther Nucleic Acids 2017;6:89–105.

6. Zhang JD, Berntenis N, Roth A, et al. Data mining reveals a network of early-response genes as a consensus signature of drug-induced in vitro and in vivo toxicity. Pharmacogenomics J 2014;14:208–216.

7. Mueller H, Wildum S, Luangsay S, et al. A novel orally available small molecule that inhibits hepatitis B virus expression. Journal of Hepatology 2018;68:412–420.

8. Drawnel FM, Zhang JD, Küng E, et al. Molecular Phenotyping Combines Molecular Information, Biological Relevance, and Patient Data to Improve Productivity of Early Drug Discovery. Cell Chem Biol 2017;24:624–634.e3.

9. Thommen DS, Koelzer VH, Herzig P, et al. A transcriptionally and functionally distinct PD-1+ CD8+ T cell pool with predictive potential in non-small-cell lung cancer treated with PD-1 blockade. Nat Med 2018;24:994–1004.

10. Rung J, Brazma A. Reuse of public genome-wide gene expression data. Nat Rev Genet 2013;14:89–99.

11. Baker M. Biotech giant publishes failures to confirm high-profile science. Nature 2016;530:141.

12. Chen C, Grennan K, Badner J, et al. Removing batch effects in analysis of expression microarray data: an evaluation of six batch adjustment methods. PLoS One 2011;6:e17238.

13. Toker L, Feng M, Pavlidis P. Whose sample is it anyway? Widespread misannotation of samples in transcriptomics studies. F1000Res 2016;5:2103.

14. Zhang JD, Hatje K, Sturm G, et al. Detect tissue heterogeneity in gene expression data with BioQC. BMC Genomics 2017;18:277.

15. Nieuwenhuis TO, Yang SY, Verma RX, et al. Consistent RNA sequencing contamination in GTEx and other data sets. Nat Commun 2020;11:1933.

16. Yoshihara K, Shahmoradgoli M, Martínez E, et al. Inferring tumour purity and stromal and immune cell admixture from expression data. Nat Commun 2013;4:2612.

17. Sturm G, Finotello F, Petitprez F, et al. Comprehensive evaluation of transcriptome-based cell-type quantification methods for immuno-oncology. Bioinformatics 2019;35:i436–i445.

18. Edgar R, Domrachev M, Lash AE. Gene Expression Omnibus: NCBI gene expression and hybridization array data repository. Nucleic Acids Res 2002;30:207–210.

19. Lachmann A, Torre D, Keenan AB, et al. Massive mining of publicly available RNA-seq data from human and mouse. Nat Commun 2018;9:1366.

20. GTEx Consortium. The Genotype-Tissue Expression (GTEx) project. Nat Genet 2013;45:580–585.

21. Lattin JE, Schroder K, Su AI, et al. Expression analysis of G Protein-Coupled Receptors in mouse macrophages. Immunome Res 2008;4:5.

22. Zhu Y, Davis S, Stephens R, et al. GEOmetadb: powerful alternative search engine for the Gene Expression Omnibus. Bioinformatics 2008;24:2798–2800.

23. Davis S, Meltzer PS. GEOquery: a bridge between the Gene Expression Omnibus (GEO) and BioConductor. Bioinformatics 2007;23:1846–1847.

24. Xie Y. bookdown. Epub ahead of print 2016. DOI: 10.1201/9781315204963.

25. Koster J, Rahmann S. Snakemake--a scalable bioinformatics workflow engine. Bioinformatics 2012;28:2520–2522.

