## Supplementary Information for "Tissue heterogeneity is prevalent in gene expression studies"

### Tissue heterogeneity is prevalent in gene expression studies (Supplementary Information)

2020-11-30

**5 References** **12**

### 1 About this document

This document is supplementary information for

> Gregor Sturm, Markus List and Jitao David Zhang. Tissue heterogeneity is prevalent in gene expression studies.

This document demonstrates step-by-step how we generated and validated tissue signatures, curated the input data, and derive our results.

The source code and instructions how to reproduce this study are available from https://github.com/grst/bioqc_geo.

### 2 Validating Tissue Signatures

The authors of *BioQC* have taken three independent approaches to show that their signatures are valid and biologically meaningful (Zhang et al. 2017). (1) They checked the results for batch effects using surrogate variable analysis, (2) they ensured that the signatures are biologically meaningful by relating them to biological knowledge, and (3) used an independent method to derive the signatures, which yields comparable results.

However, they do not quantify the performance of the signatures using standardized performance measures, such as sensitivity and specificity. In principle, one can think of three methods to achieve such a validation (Gönen 2009): (1) internal validation, *i.e.* using the same data for generating and testing signatures, (2) split-sample validation, *i.e.* dividing the dataset in a test and training dataset and (3) independent-sample validation, *i.e.* using entirely unrelated samples for training and testing. While method (2) might be acceptable if sufficient data is unavailable, only method (3) can ensure that the signature does not reflect experimental biases.

To address this, we independently derived signatures on the GTEx (Lonsdale et al. 2013) dataset using *gini-index* as described previously (Zhang et al. 2017), and performed both a 10-fold cross validation on the same dataset and a cross-species, cross-platform validation on the mouseGNF GeneAtlas (Lattin et al. 2008). To this end, we developed the python package pygenesig,[1] a framework to create and validate signatures.

In this chapter, we

- perform a 10-fold cross-validation on the GTEx dataset, calculating the precision and recall for each signature,
- perform a cross-species, cross-platform validation of the signatures generated on the GTEx dataset, and
- identify a set of tissues, that can be reliably and unabmigously identified with the *BioQC* method.

#### 2.1 Data

- The **Genotype Tissue Expression (GTEx)** project (Lonsdale et al. 2013) is a comprehensive resource of tissue-specific gene expression data. We use this dataset to derive tissue-specific signatures. The data is human only and was generated using Illumina sequencing.
- We use the **GNF Mouse GeneAtlas V3** (Lattin et al. 2008) as a control dataset to demonstrate that the gini-method is robust over multiple platforms and species. This dataset originates from mouse and was generated using the *Affymetrix Mouse Genome 430 2.0 Array (GPL1261)*.

#### 2.2 Cross-Validation of signatures on the GTEx dataset

We use *pygenesig* to create and validate signatures on the GTEx v6 dataset. The data preparation steps are performed using these jupyter notebooks.[2] The output of *pygenesig* can be viewed here.[3] Below, we summarize the analyses described in these documents.

We obtained the gene expression data and sample annotation from the GTEx portal. We collapsed gene expression data by HGNC symbol, aggregating by the sum. We aggregated samples of the same tissue by median. We generated signatures based on gini index[4] as described in the BioQC paper (Zhang et al. 2017). In brief, we calculated gini index for each gene across all tissues. Genes with a gini index $\geq 0.8$ and expression of $\geq 5$ TPM were added to the signatures of the 3 tissues with their highest expression.

[1] https://github.com/grst/pygenesig
[2] https://github.com/grst/pygenesig-example/tree/d88e4a81a45e192527a84a4445094604deba580b/notebooks/prepare_data
[3] https://github.com/grst/BioQC_GEO_analysis/blob/aa0fcd86bbdfbd49c9a4a10ce0be1c22895cc957/notebooks/gtex_v6_gini.ipynb
[4] https://grst.github.io/pygenesig/apidoc.html#module-pygenesig.gini

Next, we performed a 10-fold cross-validation as follows: We split samples in 10 stratified folds,[5] *i.e.* samples from all tissues are equally distributed across all folds. We use 9 folds to generate signatures. These signatures were applied to the remaining fold using BioQC. We iterated over the folds such that each fold has been used for training and testing.

Figure 1 shows the average BioQC score over all folds for each signature and each tissue.

As identifying contaminated or mislabled samples can be boiled down to a classification problem, we are interested in the predictive performance of each signature. Figure 2 shows the confusion matrix of using the signatures for classification. A sample is considered as classified as a tissue, if the corresponding signature scores highest among all other signatures.

#### 2.3 Reference Signatures

From the above matrices we learn that, while the vast majority of signatures yield a high score in the corresponding tissue, an unambiguous classification of tissues is only viable for a subset of tissues. For instance, the different brain regions are hard to distinguish, and so are physiologically close tissues (e.g. large and small intestine).

Here, we reduce the dataset to a subset of tissues, which can be unambigously distinguished using the BioQC method (*i.e.* precision = recall = 1.0).

We manually map[6] the tissues from GTEx to a reduced subset of tissue names. The results are available in this jupyter notebook[7] and summarized below.

Figure 3 shows the confusion matrix of the reduced signature set. All tissues have been correctly identified at precision = recall = 1.0.

#### 2.4 Cross-Platform Cross-Species Validation

To asses if the signatures translate across species and platforms, we tested the signatures generated above (human, Illumina sequencing) on the *mouseGNF tissue expression atlas* (mouse, Affymetrix microarray). The procedure is described in this notebook[8] and summarized in this section.

Figure 4 shows the score matrix of GTEx signatures against mouseGNF samples.

The signatures *Brain, Heart, Kidney, Liver, Skeletal Muscle, Pancreas, Skin* and *Testis* identify the respective tissue despite the species and platform differences at a high (>5) BioQC score.

As expected *Heart* and *Skeletal muscle* also identify each other, however *Heart* scores are still higher on heart samples and *Skeletal muscle* scores are higher on skeletal muscles samples, therefore we retain both signatures.

Surprisingly, *Adrenal Gland*, *Ovary* and *Uterus* are not able to identify the respective samples, despite having a high score in the cross-validation. We, therefore, exclude these signatures from the reference signature set.

Unfortunately, *Blood* was not profiled in the mouseGNF dataset. We keep the signature nonetheless as it does not trigger any false positives.

[5] http://scikit-learn.org/stable/modules/generated/sklearn.model_selection.StratifiedKFold.html

[6] https://github.com/grst/pygenesig-example/blob/d88e4a81a45e192527a84a4445094604deba580b/manual_annotation/gtex_solid.csv

[7] https://github.com/grst/BioQC_GEO_analysis/blob/b11987da13ba9b98eba34206f92942be8de6427e/signature_validation/gtex_v6_gini_solid.ipynb

[8] https://github.com/grst/pygenesig-example/blob/80bfe2a388a5230b004c288cb2ea220f0394855d/notebooks/gtex_solid_vs_mouse_gnf.ipynb

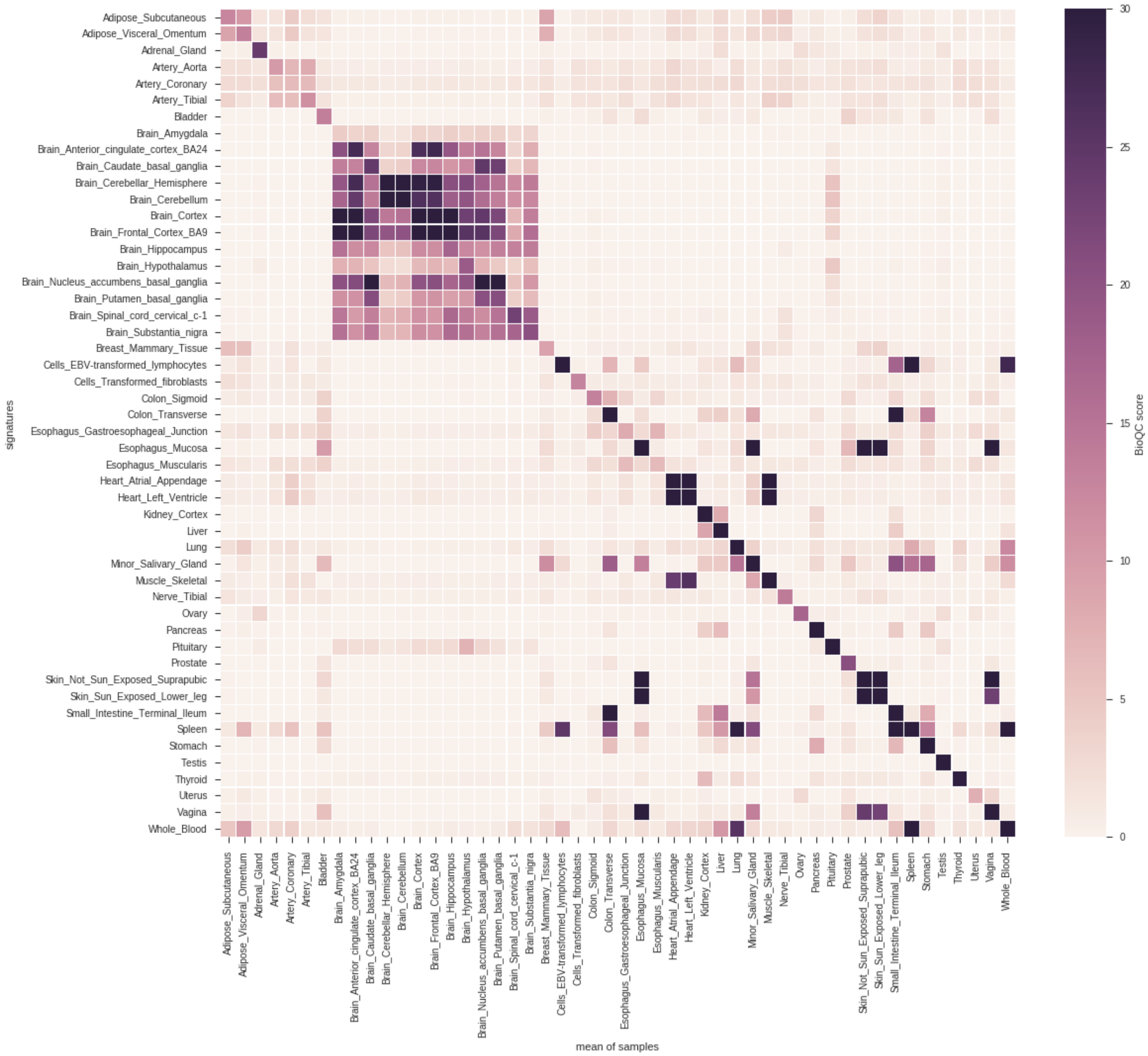

Figure 1: cross-validation of GTEx tissue signatures. Signatures are shown on the y-axis, the corresponding groups of samples on the x-axis. The tile shading indicates the average BioQC score of a signature on a group of samples. For better visibility of low scores, the colors are saturated at 30.

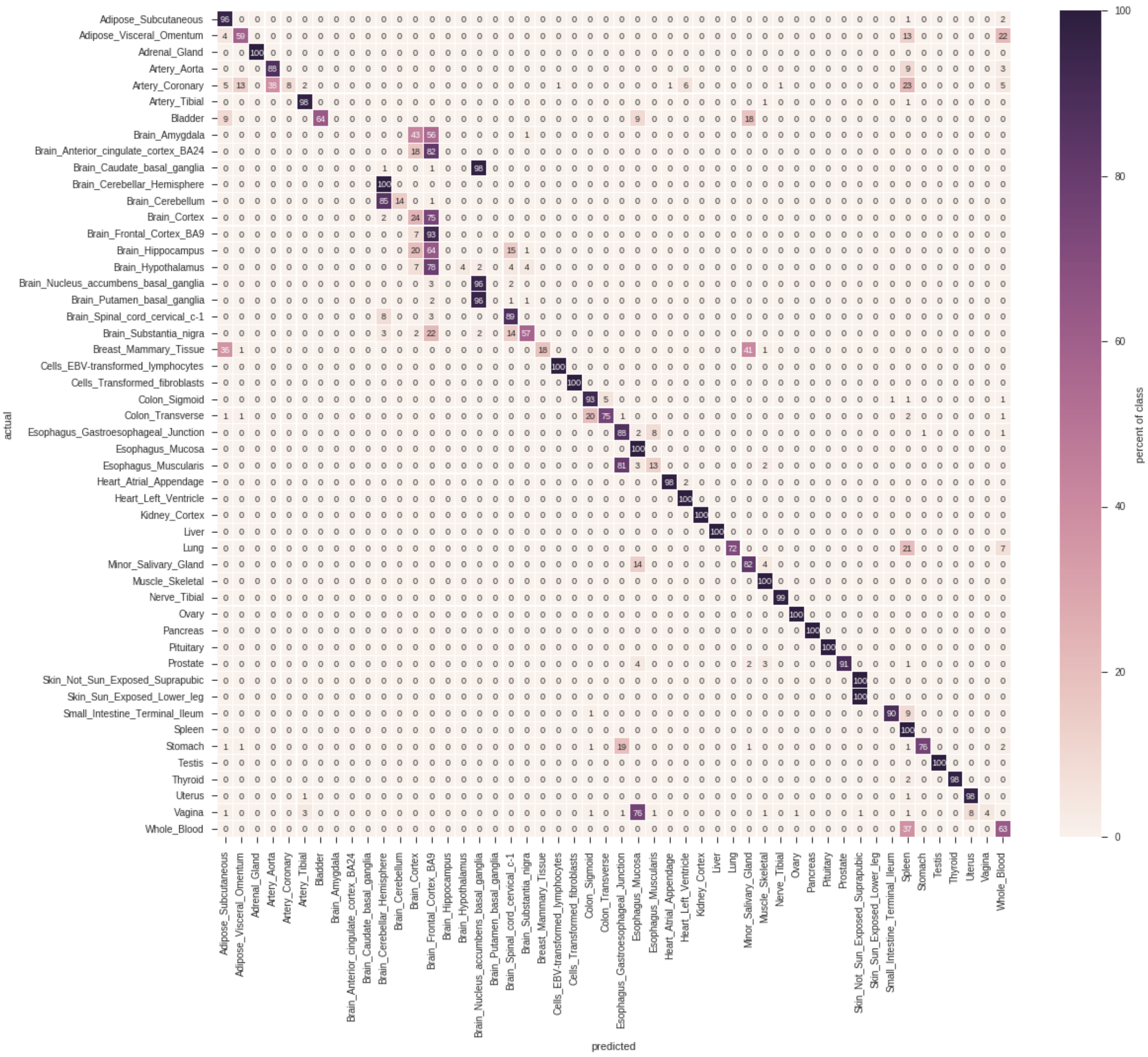

Figure 2: Confusion matrix of the cross-validation.

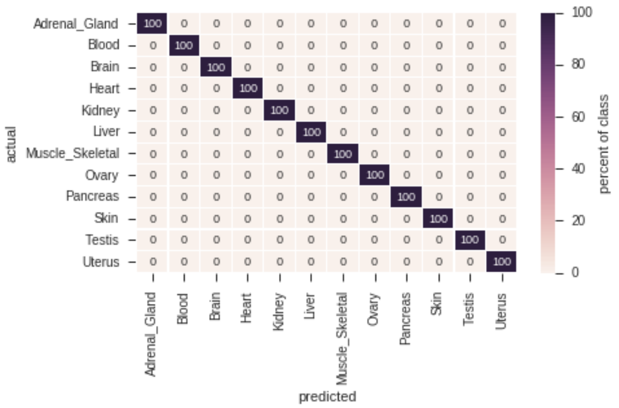

Figure 3: Confusion matrix of robust signatures.

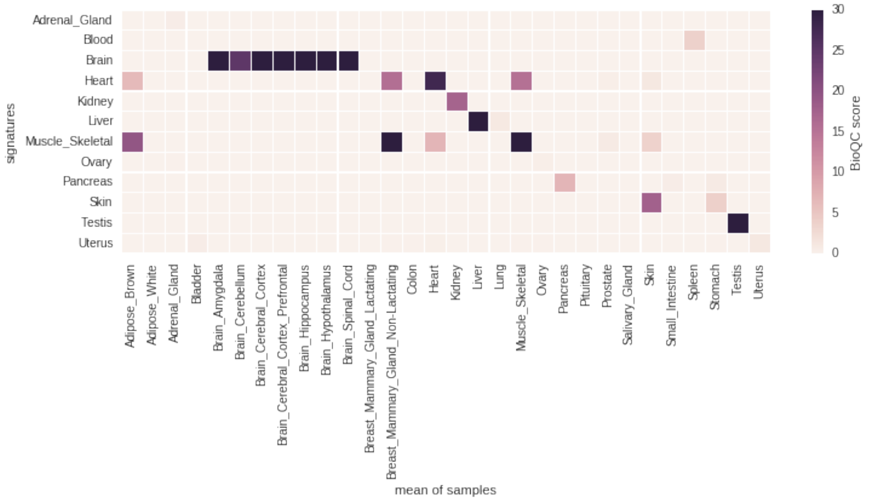

Figure 4: Cross-platform, cross-species validation of the robust signatures identified in the previous step on mouse microarray data.

### 3 Sample data and metadata

In this section, we describe how we obtained and curated gene expression data and sample metadata.

#### 3.1 Standardize Tissue Names

The annotation of tissues is inconsistent within GEO and ARCHS4. A "liver" sample can be termed *e.g.* "liver", "liver biopsy" or "primary liver". We, therefore, need a way to *standardize* the tissue name. We manually mapped the most abundant tissues to a controlled vocabulary. Next, in order to find out which samples are subject to *tissue heterogeniety*, we first need to define which signatures we would expect in a certain tissue. For example, we map the signatures `Intestine_Colon_cecum_NR_0.7_3` and `Intestine_Colon_NR_0.7_3` to *colon*. Since the reference signatures are not as specific as the tissue annotation, we created *tissue sets* to combine them into groups. For instance, it is hard to distinguish *jejunum* from *colon*, but easy to distinguish the two from other tissues. We therefore created a tissue set *intestine*, which contains both *jejunum* and *colon* and references all signatures associated with the two tissues. All of the described mappings are available from Supplementary Table 1.[9]

#### 3.2 GEO

##### 3.2.1 Downloading GEO data

We retrieved sample metadata for GEO using the GEOmetadb package (Zhu et al. 2008). We downloaded the studies with GEOquery (Davis and Meltzer 2007) and stored them as R ExpressionSet (Huber et al. 2015) using the R script geo_to_eset.R.[10] We used the `annotGPL=TRUE` option of GEOquery's `getGEO` function to obtain gene symbols for the studies, where available. Since the tissue signatures use human gene symbols, we added human orthologs for all mouse and rat samples.

[9] https://github.com/grst/BioQC_GEO_analysis/blob/master/manual_annotation/tissue_annotation.xlsx
[10] https://github.com/grst/BioQC_GEO_analysis/blob/master/scripts/geo_to_eset.R

##### 3.2.2 Filtering GEO data

We filtered GEO samples by the following criteria (figure 5):

1. the tissue or origin is annotated,
2. gene symbols are annotated,
3. the readout was performed on a single-channel microarray, and
4. the tissue could be mapped to the controlled vocabulary (section 3.1).
5. We only retained samples from the three major organisms: human, rat, mouse.
6. We removed studies which have been normalized per-gene and where ubiquitous house-keeping genes were not expressed.
7. Finally, we only retained samples originating from tissues for which a *reference signature* is available.

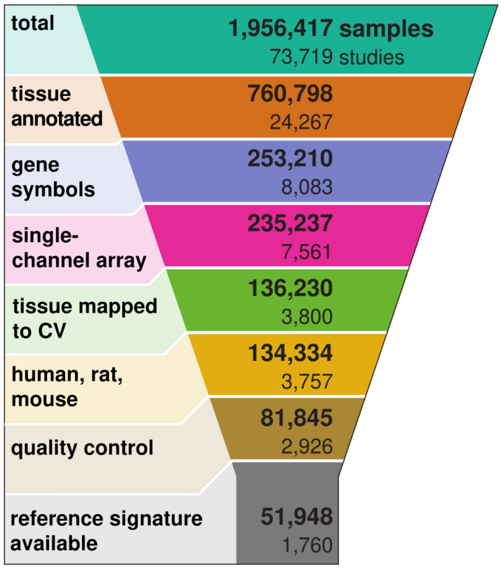

Figure 5: Summary of filtering steps on GEO samples

#### 3.3 ARCHS4

In addition to GEO, we used data from ARCHS4 (Lachmann et al. 2018), a publicly available data collection of annotaed, consistently processed gene expression datasets based on RNA-sequencing. We downloaded gene expression and metadata as `RData` objects from the ARCHS4 website (version 8.0).[11]

We filtered samples by the following criteria (figure 6):

1. The library is a transcriptomic cDNA library, the library strategy is RNA-seq, and either polyA or total RNA were extracted.
2. No single-cell RNA-seq samples (none of the annotation fields may contain the keywords "single-cell", "single cell" or "smartseq")
3. At least 500,000 reads could be mapped to genes.
4. The tissue could be mapped to the controlled vocabulary (section 3.1).
5. Finally, we only retained samples originating from tissues for which a *reference signature* is available.

Gene counts were normalized into TPM values before analysing them with BioQC.

[11] https://amp.pharm.mssm.edu/archs4/download.html

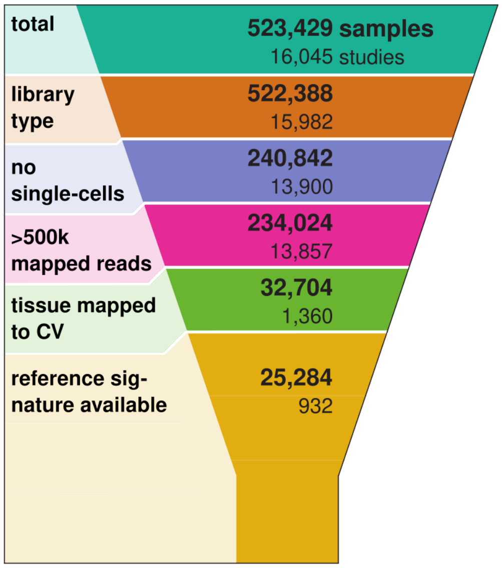

Figure 6: Summary of filtering steps on ARCHS4 samples

Table 1: reference signatures

| Reference Signature | Tissue |
|---|---|
| GTEX_Blood | blood |
| GTEX_Brain | brain |
| GTEX_Heart | heart |
| GTEX_Kidney | kidney |
| GTEX_Liver | liver |
| GTEX_Muscle_Skeletal | skeletal muscle |
| GTEX_Pancreas | pancreas |
| GTEX_Skin | skin |
| GTEX_Testis | testis |

### 4 Testing for tissue heterogeneity

#### 4.1 Tissue signatures

In section 2, we identified a set of 9 *reference signatures* (table 1) which unambiguously identify their corresponding tissue across platforms and species. In addition to that, we use 120 tissue signatures from the BioQC publication, which we refer to as *query signatures*.

#### 4.2 Testing samples for heterogeneity

We tested for enrichment of 120 selected signatures provided by BioQC and the 9 reference signatures generated by us on all 76576 selected samples resulting in a list of 10031456 (sample, signature, pvalue) pairs.

Our intention is to identify samples that show *tissue heterogeneity*, *i.e.* unintentional profiling of cells of other origin than the target tissue of profiling. We classify samples into *heterogeneous* and *not heterogeneous*. We call a classification *true-positive* if the given sample is classified as *heterogeneous* and the sample indeed contains cells different from the annotated tissues. Analogous, we call a classification *false-positive* if the given sample is classified as *heterogeneous* but in reality only contains cells from the annotated tissue.

Naively, we could label a sample as heterogeneous, if a signature unrelated to the annotated tissue exceeds a certain score. The problem with this approach is, that some signatures overlap; the resulting scores are therefore correlated and will lead to false-positives. One cannot simply solve this problem by excuding genes that are members of multiple signatures, as it is easily possible to build two (in fact many) distinct, non-overlapping signatures matching the same tissue.

In section 2 we have created and validated *reference signatures* for 9 tissues. Even though we have demonstrated that each signature unambigously identifies its corresponding tissue (*i.e.* scores highest), the signatures could still be correlated. Some of them in fact are, e.g. cardiac muscle and skeletal muscle (see figure 7). Moreover, we lack sufficient data to perform an independent-sample validation on the signatures provided by BioQC. Accounting for correlation with the reference signatures, we can still avoid false-positives, even if a signature is characteristic for a tissue histologically close to the annotated tissue.

We, therefore, take the following approach to account for the correlation of signatures: A given sample $s$ annotated as tissue $t$ is tested for enrichment with signature $k_{\text{query}}$ resulting in a p-value $p_{\text{query}}$. Let $k_{\text{ref}}$ be the reference signature associated with tissue $t$ and $p_{\text{ref}}$ the p-value of testing $s$ for enrichment of $k_{\text{ref}}$. Let $\tau$ be a certain false discovery rate (FDR)-threshold.

(1) If the Benjamini-Hochberg (BH)-adjusted $p_{\text{query}} \geq \tau$, we assume that $s$ is not heterogeneous; else continue.
(2) We fit a linear model using `rlm` from the `R` `MASS` package of $|log10(p_{\text{query}})|$ against $|log10(p_{\text{ref}})|$ for all samples annotated as $t$. We assume that the residuals $R$ of the linear model follow a normal distribution $R \sim \mathcal{N}(0, \sigma^2)$, where $\sigma$ is the standard deviation of the residuals.
(3) We extract the residual $r$ corresponding to sample $s$. We calculate the p-value $p_{\text{corr}} = 1 - \text{CDF}_R(r)$ where $\text{CDF}_R$ is the cumulative density function of $\mathcal{N}(0, \sigma^2)$.
(4) If the BH-adjusted $p_{\text{corr}} < \tau$, we reject the hypothesis that $k_{\text{query}}$ is enriched only due to correlation and label the sample as heterogeneous.

###### 4.2.1 Determining $\tau$.

We desire an overall FDR of 0.01. Since we test in a two-step procedure, we choose a threshold $\tau$, such that the overall FDR equals 0.01.

$$\text{FDR} = \tau + (1 - \tau)\tau$$

Explanation: Of all positives in step 1, there are, per definition, a fraction of $\tau$ false-positives and $1 - \tau$ true-positives. To all positives, the step 2 test is applied. In addition to the false-positives from step 1, there is a fraction of $\tau(1 - \tau)$ false-positives among the true-positives from step 1.

Solving the equation yields:

$$0.01 = \tau + (1 - \tau)\tau \Leftrightarrow 0 = \tau^2 - 2\tau + 0.01$$

```
tau = polyroot(c(FDR_THRES, -2, 1))[1]
tau
```

```
## [1] 0.005012563-0i
```

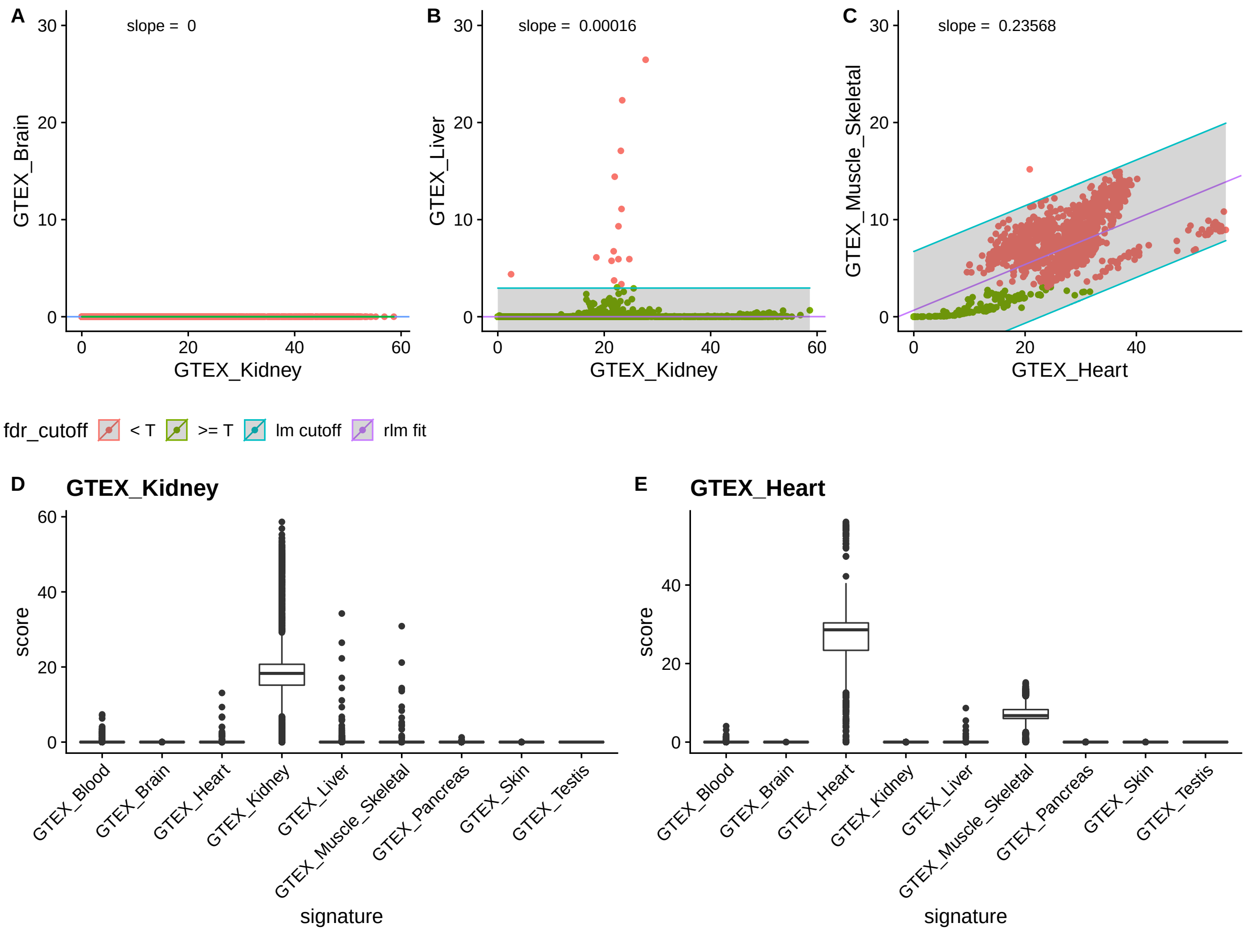

Figure 7: Examples of signature correlation. Panels A-C: correlation of the signature scores (y-axis) against the scores of a reference signature (x-axis). The purple line indicates the model fitted to the data. Points falling into the shaded area can be explained by the linear model. Points are colored according to whether they would be discarded already in the FDR-filtering step. (A) Brain scores of kidney samples against the scores of the kidney signature. There is virtually no correlation, nor are there outliers. Apparently, no kidney samples are contaminated with neuronal cells. (B) Liver scores of kidney samples against scores of the kidney signature. The samples are not correlated, however some outliers are detected which are samples potentially containing liver cells. (C) Cardiac muscle scores of skeletal muscle samples against skeletal muscle scores. The scores are highly correlated. While most of the points exceed the FDR threshold, they will not be classified as outliers because the elevated score can be explained by the correlation. Panels D and E show the boxplots of the scores of various signatures on kidney and heart samples, respectively.

### 5 References

Davis, S., and P. S. Meltzer. 2007. "GEOquery: A Bridge Between the Gene Expression Omnibus (GEO) and BioConductor." *Bioinformatics* 23 (14): 1846–7. https://doi.org/10.1093/bioinformatics/btm254.

Gönen, Mithat. 2009. "Statistical aspects of gene signatures and molecular targets." *Gastrointest Cancer Res* 3 (2 Suppl): 19–21. http://www.pubmedcentral.nih.gov/articlerender.fcgi?artid=2684735%7B/&%7Dtool=pmcentrez%7B/&%7Drendertype=abstract.

Huber, Wolfgang, Vincent J Carey, Robert Gentleman, Simon Anders, Marc Carlson, Benilton S Carvalho, Hector Corrada Bravo, et al. 2015. "Orchestrating High-Throughput Genomic Analysis with Bioconductor." *Nature Methods* 12 (2): 115–21. https://doi.org/10.1038/nmeth.3252.

Lachmann, Alexander, Denis Torre, Alexandra B. Keenan, Kathleen M. Jagodnik, Hoyjin J. Lee, Lily Wang, Moshe C. Silverstein, and Avi Ma'ayan. 2018. "Massive Mining of Publicly Available RNA-Seq Data from Human and Mouse." *Nature Communications* 9 (1). https://doi.org/10.1038/s41467-018-03751-6.

Lattin, Jane E, Kate Schroder, Andrew I Su, John R Walker, Jie Zhang, Tim Wiltshire, Kaoru Saijo, et al. 2008. "Expression Analysis of G Protein-Coupled Receptors in Mouse Macrophages." *Immunome Research* 4 (1): 5. https://doi.org/10.1186/1745-7580-4-5.

Lonsdale, John, Jeffrey Thomas, Mike Salvatore, Rebecca Phillips, Edmund Lo, Saboor Shad, Richard Hasz, et al. 2013. "The Genotype-Tissue Expression (GTEx) project." *Nature Genetics* 45 (6): 580–85. https://doi.org/10.1038/ng.2653.

Zhang, Jitao David, Klas Hatje, Gregor Sturm, Clemens Broger, Martin Ebeling, Martine Burtin, Fabiola Terzi, Silvia Ines Pomposiello, and Laura Badi. 2017. "Detect tissue heterogeneity in gene expression data with BioQC." *BMC Genomics* 18 (1): 277. https://doi.org/10.1186/s12864-017-3661-2.

Zhu, Yuelin, Sean Davis, Robert Stephens, Paul S. Meltzer, and Yidong Chen. 2008. "GEOmetadb: Powerful alternative search engine for the Gene Expression Omnibus." *Bioinformatics* 24 (23): 2798–2800. https://doi.org/10.1093/bioinformatics/btn520.
